# CRISPRa-mediated activation of genes associated with inherited retinal dystrophies in acutely isolated human cells for diagnostic purposes

**DOI:** 10.1101/2024.11.29.625601

**Authors:** Valentin J. Weber, Alice Reschigna, Maximilian-J. Gerhardt, Klara S. Hinrichsmeyer, Dina Y. Otify, Thomas Heigl, Frank Blaser, Isabelle Meneau, Martin Biel, Stylianos Michalakis, Elvir Becirovic

**Affiliations:** Laboratory for Retinal Gene Therapy, Department of Ophthalmology, University Hospital Zurich, University of Zurich, Zurich, Switzerland; Department of Ophthalmology, LMU University Hospital, LMU Munich, Munich, Germany; Department of Pharmacy – Center for Drug Research, LMU Munich, Munich, Germany; Department of Ophthalmology, University Hospital Zurich, Zurich, Switzerland

**Keywords:** Diagnostics, Genetic diseases, RNA processing

## Abstract

Many patients suffering from inherited diseases do not receive a genetic diagnosis and are therefore excluded as candidates for treatments, such as gene therapies. Analyzing disease-related gene transcripts from patient cells would improve detection of mutations that have been missed or misinterpreted in terms of pathogenicity during routine genome sequencing. However, the analysis of transcripts is complicated by the fact that a biopsy of the affected tissue is often not appropriate, and many disease-associated genes are not expressed in tissues or cells that can be easily obtained from patients. Here, using CRISPR/Cas-mediated transcriptional activation (CRISPRa) we developed a robust and efficient approach to activate genes in skin-derived fibroblasts and in freshly isolated peripheral blood mononuclear cells (PBMCs) from healthy individuals. This approach was successfully applied to blood samples from patients with inherited retinal dystrophies (IRD). We were able to efficiently activate several IRD-linked genes and detect the corresponding transcripts using different diagnostically relevant methods such as RT-qPCR, RT-PCR and long- and short-read RNA sequencing. The detection and analysis of known and unknown mRNA isoforms demonstrates the potential of CRISPRa-mediated transcriptional activation in PBMCs. These results will contribute to ceasing the critical gap in the genetic diagnosis of patients with IRD or other inherited diseases.

**Graphical abstract:** 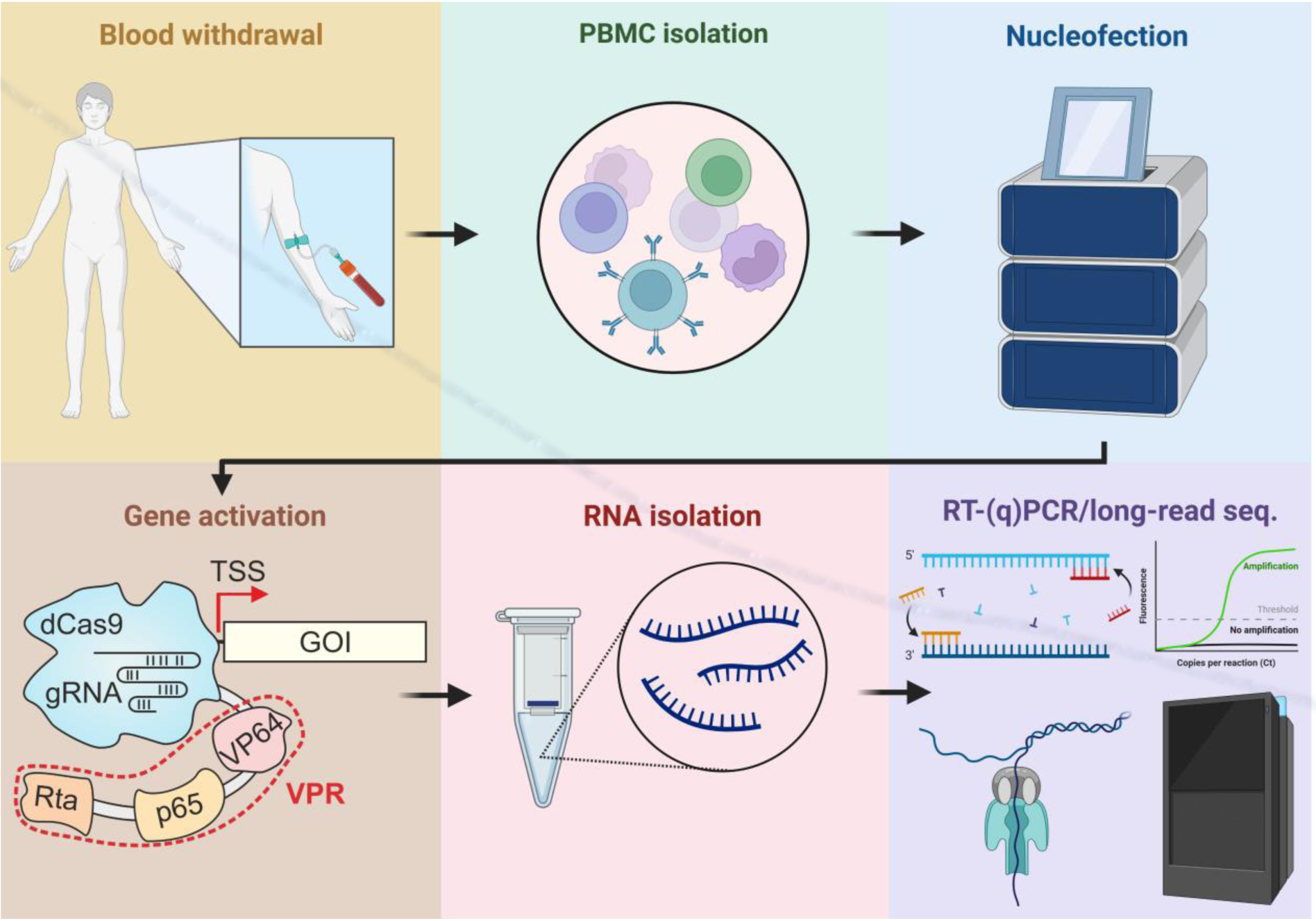

## Introduction

Even in developed countries with modern molecular diagnostic equipment, up to 50% of patients with genetic diseases receive no or insufficient genetic diagnosis (1). This excludes patients from established treatments and participation in clinical trials testing novel therapies for the respective diseases.

High-throughput next generation sequencing (NGS) technologies, such as whole genome sequencing (WGS) or whole exome sequencing (WES) have facilitated the diagnostics of genetic diseases. Both WGS and WES, however, have key limitations. WES does not cover the non-coding regions (e.g. introns, promoters or other regulatory transcriptional elements), which are crucial for mRNA stability and processing. WGS is time and money-consuming, and the interpretation of the large amount of data obtained during this process is rather difficult (2, 3). Even if potential disease-causing mutations can be identified in coding or non-coding regions of candidate genes using WGS, experimental validation of how these mutations influence the mRNA level is inevitable but laborious using currently available techniques (e.g. minigene assays (4) or patient-derived iPSC models). Single nucleotide variants can affect mRNA processing via different mechanisms. Among those, the most common one alters pre-mRNA splicing (5). Splice site mutations are typically classified as such using standard splice prediction software (6, 7). Nevertheless, bioinformatic prediction of the effects of mutations is often not reliable. Apart from false-positive results, splice prediction programs also yield an uncertain number of false-negative records. Many false-negative splicing mutations are classified as missense or silent variants located in (deep) exonic coding regions (8, 9). However, such point mutations can also affect regulatory splice elements, ultimately leading to aberrant splicing, and should therefore be routinely tested at the transcript level, preferably in patient cells (10).

Taken together, there is an unmet need for developing straightforward techniques focusing on detection and investigation of disease-associated mutations at the transcript level. The most convenient method to analyze the transcripts of the corresponding genes is to use patients’ tissue samples. However, taking biopsies of affected tissues may often not be reasonable and many disease-relevant genes are not expressed in cells which can be routinely obtained from the patients (e.g. blood cells or skin-derived fibroblasts). This applies in particular to IRDs, in which tissue collection from the affected retina for diagnostic purposes is generally not justified due to the invasiveness of the procedure, risk of iatrogenic damage and potentially sight-threatening complications. To circumvent these obstacles, we developed a <u>C</u>RISPRa-based <u>a</u>pproach for transcriptional <u>a</u>ctivation of genes in acute<u>ly</u> isola<u>te</u>d human <u>c</u>ells (hereinafter referred to as CATALYTEC) to analyze single or multiple genes associated with IRDs.

## Results

To provide proof of principle for CATALYTEC, we initially focused on *ABCA4*, *RPE65*, *MYO7A* and *USH2A* for several reasons:

i. The individual genes are frequently mutated in the corresponding diseases: Stargardt disease (STGD), Leber congenital amaurosis (LCA), Retinitis pigmentosa (RP), and Usher syndrome (USH).
ii. Three of the four genes (*USH2A*, *MYO7A*, *and ABCA4*) have a large gene body, a condition that complicates the identification of potentially pathogenic mutations, particularly those located in non-coding regions.
iii. *RPE65* offers high therapeutic relevance as *RPE65* retinopathy can be treated with the approved AAV gene supplementation therapy voretigene neparvovec. Improved genetic testing could help to confirm pathogenicity of novel *RPE65* variants of uncertain significance and thus qualify affected patients for treatment with the approved gene therapy.

Our initial gene activation experiments were performed and optimized in HEK293T cells transfected with cassettes expressing nuclease deficient dCas9-VPR, one of the most effective CRISPRa systems (11, 12), in combination with single guide (sg) RNAs targeting the transcriptional start site (TSS) region of the respective genes (Fig. 1A). We were able to identify sgRNAs for efficient activation of each of these genes. Multiplexing experiments demonstrated that the combination of multiple sgRNA cassettes resulted in simultaneous activation of all genes without any substantial loss of activation efficiency (Fig. 1D). Using RT-PCR, we could amplify all fragments covering the entire coding region and parts of the untranslated regions (Fig. 1B+C, Suppl. Fig. 1). The identity of all bands was confirmed by Sanger sequencing.

**Figure 1.**
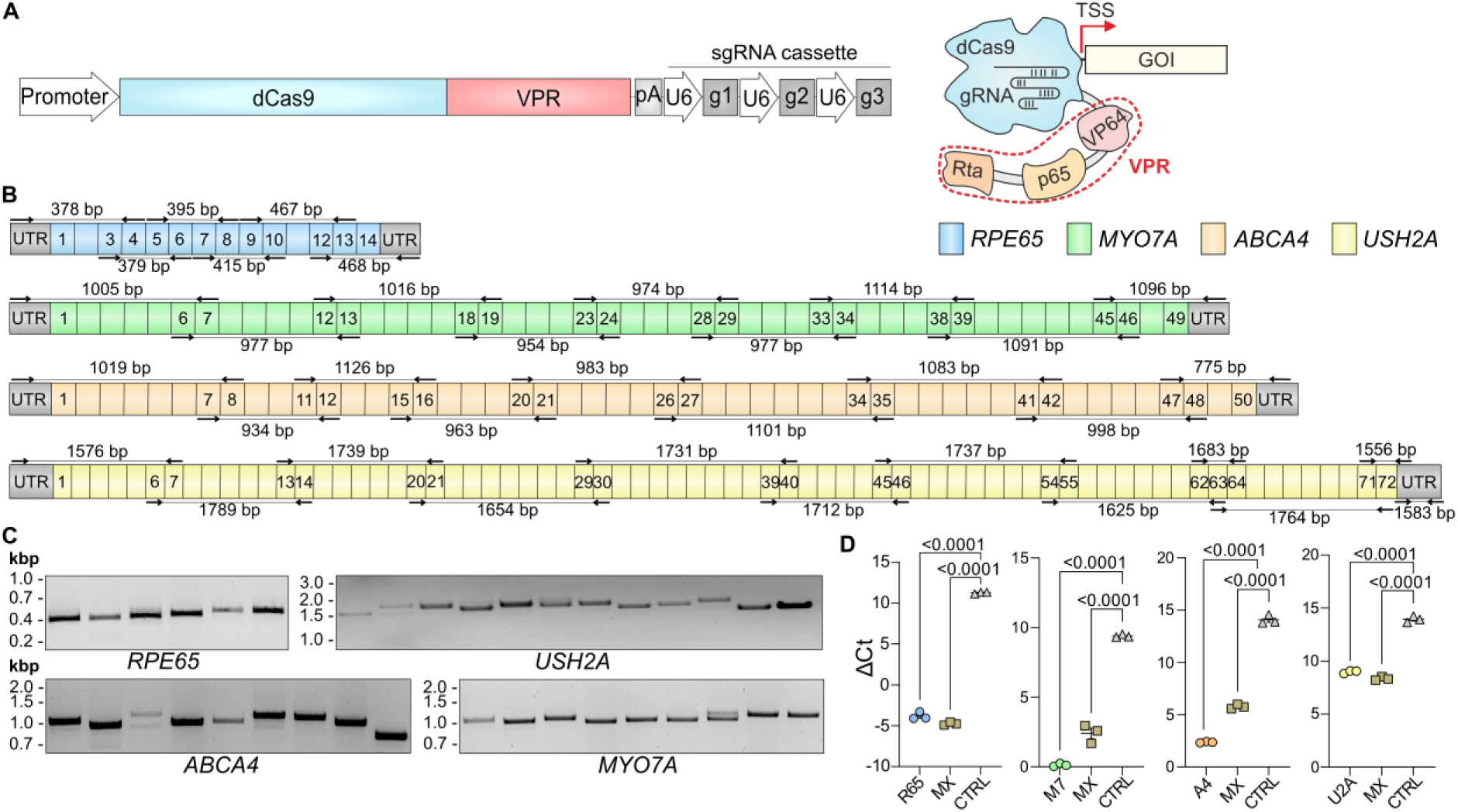
Transcriptional activation of IRD-associated genes in transfected HEK293T. **A**, Scheme depicting the expression plasmid and the corresponding dCas9-VPR protein bound to a gene of interest (GOI). dCas9-VPR, driven by either CMV or SFFV promoter, was combined with a cassette expressing sgRNAs, targeting either *RPE65*, *ABCA4*, *MYO7A* or *USH2A*. For simultaneous gene activation multiple sgRNA cassettes were combined. pA, poly A signal. **B**, Primer design for RT-PCR analysis. Primer binding sites (black arrows) and PCR product lengths in base pairs (bp) are shown in the schematic representation of the transcripts. Colored boxes represent numbered exons of the respective genes. UTR, untranslated region. **C**, RT-PCR result for each gene transcript after transcriptional activation in HEK293T cells. kbp, kilo base pairs. **D**, RT-qPCR result for each gene activation. MX, multiplexed gene transcriptional activation by combination of multiple sgRNA cassettes. R65, *RPE65*; M7, *MYO7A*; A4, *ABCA4*; U2A, *USH2A*. Values are shown as mean ± SEM. Statistics were calculated with Student’s t-test.

In the first attempts to establish a robust and simple protocol for the transfection of PBMCs and skin fibroblasts from healthy individuals, we tested various commonly used transfection techniques (e.g. calcium phosphate, lipofection and electroporation). For most of these approaches, however, we observed no or weak expression of the transfected GFP reporter gene.

Since lentiviral vectors (LV) have been shown to efficiently transduce non-dividing cells (13), we investigated the efficacy of gene activation upon transduction with LVs. In our experiments, using dual LV vectors, each expressing GFP or dsRed, we obtained high co-transduction efficiencies for human skin fibroblasts and PBMCs (Suppl. Fig. 2A, B). Surprisingly, the dual LV approach for gene transcriptional activation in transduced HEK293T, fibroblasts and PBMCs led to a relatively weak transcriptional activation of *ABCA4*, despite numerous optimization steps (e.g. use of high multiplicity of infections (MOIs), different promoters etc., Suppl. Fig. 2C-F). These results suggest that dCas9-VPR delivered via LVs is substantially less efficient in activating target genes compared to delivery by plasmid transfection. To elucidate the causes for the low activity of dCas9-VPR in cells transduced with the corresponding LVs, we generated a HEK293T cell line stably expressing dCas9-VPR through genomic integration of the corresponding cassette (Suppl. Fig. 3A). After transduction of these cells with LVs expressing only the sgRNAs targeting the individual genes, we detected a robust transcriptional activation of *ABCA4*, *MYO7A* and *USH2A*. A similarly high activation was also achieved in a multiplexing approach, in which all three genes were targeted simultaneously (Suppl. Fig. 3B). In combination with the results shown in Suppl. Fig. 2, this indicates that the efficiency of transcriptional activation is not compromised by LV-mediated delivery of sgRNAs, excluding a general impact of LVs on gene activation. A possible explanation for our observation of low-level gene activation in all cell types upon LV-based administration of dCas9-VPR is that dCas9-VPR is not well tolerated by LV vectors due to its size (5.8 kb). To test this hypothesis, we used a shorter variant of this CRISPRa system, dCas9-VPRmini (14) (Suppl. Fig. 3C). However, this approach also did not lead to increased activation efficiency compared to full-length dCas9-VPR (Suppl. Fig. 3D).

From this set of unfavorable results, we concluded that the LV-based approach in combination with dCas9-VPR mediated transcriptional activation of genes is unsuitable for diagnostic purposes.

Achieving reasonable levels of transcriptional activation in PBMCs or skin fibroblasts from healthy individuals is a prerequisite for establishing a robust protocol for diagnostic purposes in patient cells. So far, although we initially obtained high activation efficiencies for the individual IRD-linked genes in transfected HEK293T cells, none of the previously described approaches has reached this goal. In another attempt to achieve this milestone, we used nucleofection for transferring sgRNA and transgene plasmids into our primary target cells. Nucleofection of a GFP-encoding cassette resulted in robust expression of this reporter in both cell types (Suppl. Fig. 4A). Next, we nucleofected the dCas9-VPR expression cassette together with sgRNAs targeting either *MYO7A*, *USH2A*, *ABCA4* or *RPE65*. Using RT-qPCR, we observed high activation efficiencies for *ABCA4* and *RPE65* in both cell types. Additionally, we were able to detect the entire coding and untranslated regions using RT-PCR (Fig. 2A, B and Suppl. Fig. 5). By comparison, we found that *MYO7A* is already endogenously expressed in human skin fibroblasts and in PBMCs, respectively, in sufficient amounts to be detected by RT-PCR. Nevertheless, after transcriptional activation a moderate increase in *MYO7A* expression was detectable via RT-qPCR (Suppl. Fig. 4B, C). For *USH2A*, we could only detect parts covering exons 1-7, 45-55 and the 3’-UTR of the corresponding mRNA (Suppl. Fig. 4D, E).

**Figure 2.**
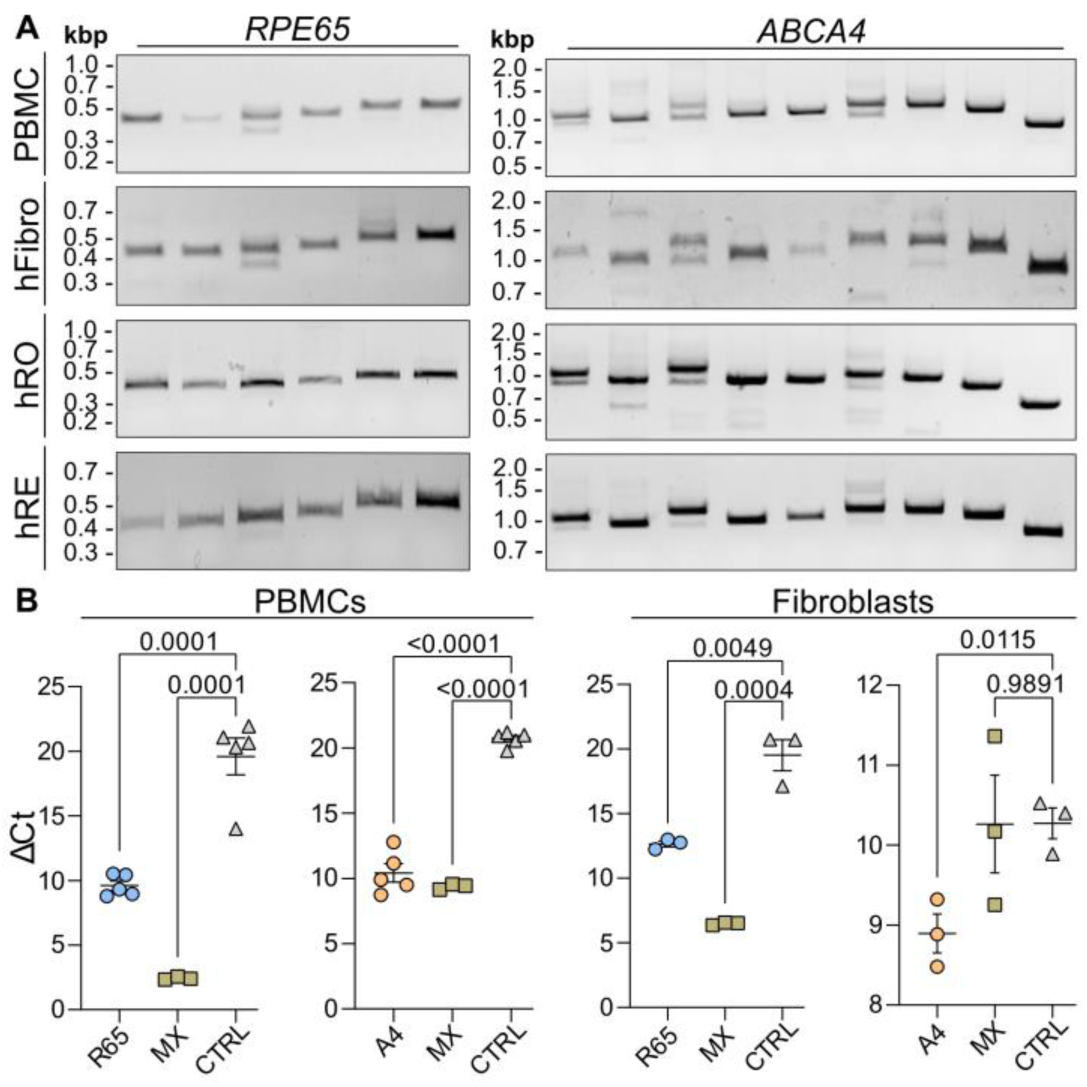
RT-(q)PCR analyses of transcriptionally activated genes in human PBMCs and fibroblasts. **A**, RT-PCR results for *RPE65* (left) *and ABCA4* (right). Transcriptionally activated PBMCs and fibroblasts are compared to endogenously expressed *RPE65* and *ABCA4* from human retinal organoids (hROs) and human retina (hRE). **B**, RT-qPCR results for transcriptionally activated *RPE65* (R65) and *ABCA4* (A4) in PBMCs (left) and fibroblasts (right). MX, multiplexed transcriptional activation of both genes. Values are given as mean ± SEM. Statistics are calculated with Student’s t-test.

For *RPE65* and *ABCA4,* in addition to the expected RT-PCR bands encoding the full-length proteins, we detected several splice variants, most of which resulted in frameshifts and premature termination codons (PTC, Suppl. Tab. 1 + 2). The corresponding bands could represent alternative splicing that might differ in PBMCs and in the retina.

To test how the splice pattern observed in PBMCs compared to that of human retinal cells, we performed RT-PCR of RNA samples isolated from human donor retinas and human retinal organoids using the same primer pairs. We observed a very similar band pattern compared to the RT-PCR results from skin fibroblasts and PBMCs, suggesting no major differences in mRNA splicing of *ABCA4* and *RPE65* between the different cell and tissue types (Fig. 2A).

Although RT-PCR is a robust method for the qualitative detection of mRNA splicing defects, it is less suitable for the reliable quantification of the results. In addition, particularly for low abundant transcripts, it is susceptible to artificial variations, manifesting during cDNA synthesis and subsequent PCR amplification. To overcome the drawbacks of RT-PCR and to validate our results with additional methods, we analyzed RNA from PBMCs of several healthy individuals using two next generation RNA sequencing techniques: short-read RNA sequencing (Illumina) and long-read RNA sequencing (PacBio Revio, Fig. 3A). Short-read sequencing of samples with transcriptionally activated *ABCA4* and *RPE65* showed a robust increase in gene expression for both genes in comparison to non-treated control samples (Fig. 3B). Splice junction analysis of *ABCA4* demonstrated that the level of transcriptional activation achieved with CATALYTEC is in principle sufficient to analyze the different transcript isoforms (Suppl. Fig. 4F).

**Figure 3.**
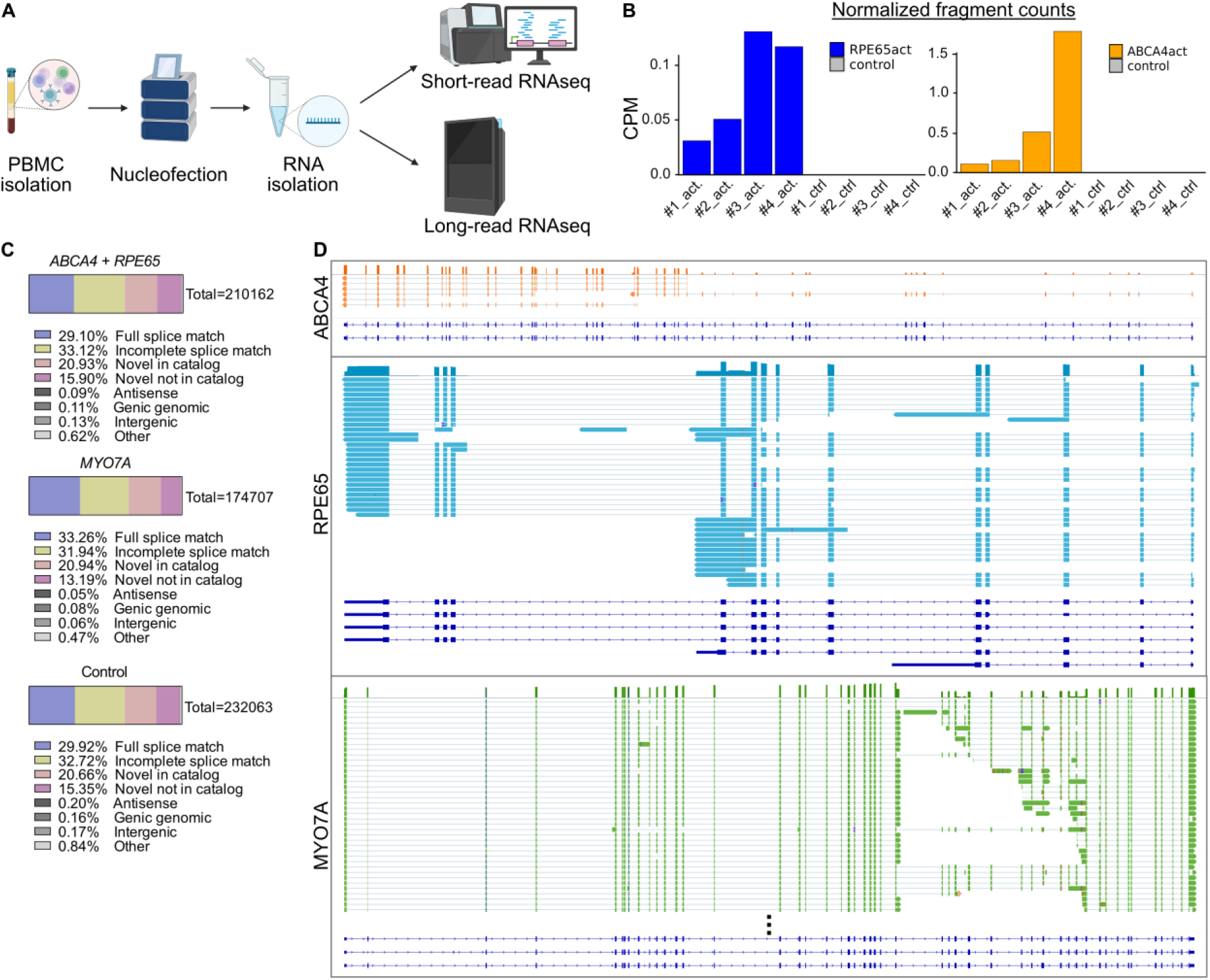
Short- and long-read RNA sequencing of transcriptionally activated PBMCs. **A**, Workflow diagram from PBMC isolation to short- and long-read RNA sequencing. **B**, Normalized fragment counts for *RPE65* (left, blue) and *ABCA4* (right, orange) of four analyzed samples (#1-4) compared to untreated control samples. CPM, counts per million. **C**, Proportion of different structural variant categories after long-read sequencing, clustering and classification. Total number represents the value of 100% for each sample. **D**, Total transcripts covering reads for either *ABCA4* (orange), *RPE65* (blue) or *MYO7A* (green) after long-read sequencing. Dark blue schemes below represent the reference transcripts of the respective genes.

Analysis of long-read RNA sequencing data in healthy PBMC samples revealed that over 30% of detected splice variants exhibit incomplete splice matches, i.e. variants with partial alignment but differing at the 5’ or/and 3’-end of the transcript. Additionally, more than 30% of isoforms represent either novel combinations of known splice sites (classified as “Novel in catalogue”) or incorporate at least one previously unidentified splice site (“Novel not in catalogue”), highlighting the capability of long-read sequencing to identify novel splice isoforms (Fig. 3C).

Simultaneous transactivation of *ABCA4* and *RPE65* and single transactivation of *MYO7A* led to full transcript coverage by long-read sequencing, corroborating the increase in gene expression as previously determined via RT-qPCR and short-read RNA sequencing (Fig. 3D). The respective consensus sequences, constructed by clustering similar reads and eliminating potential artifacts, were used to identify isoforms of biological relevance. For *RPE65* and *MYO7A,* several full-length isoforms were identified as consensus sequences (Suppl. Fig. 6). In contrast, only one consensus sequence was obtained for *ABCA4*, as multiple transcript reads were excluded in the filtering process.

Finally, we applied the described CATALYTEC protocol to one patient with confirmed biallelic *RPE65*-associated LCA (P1), two patients with the clinical phenotype of rod-cone dystrophy (RP, P2 and P3) but unclear molecular genetic diagnosis, and three patients with the clinical diagnosis of *ABCA4*-associated Stargardt’s disease (STGD1; P4-P6) (Tab. 1, Fig. 4).

**Figure 4.**
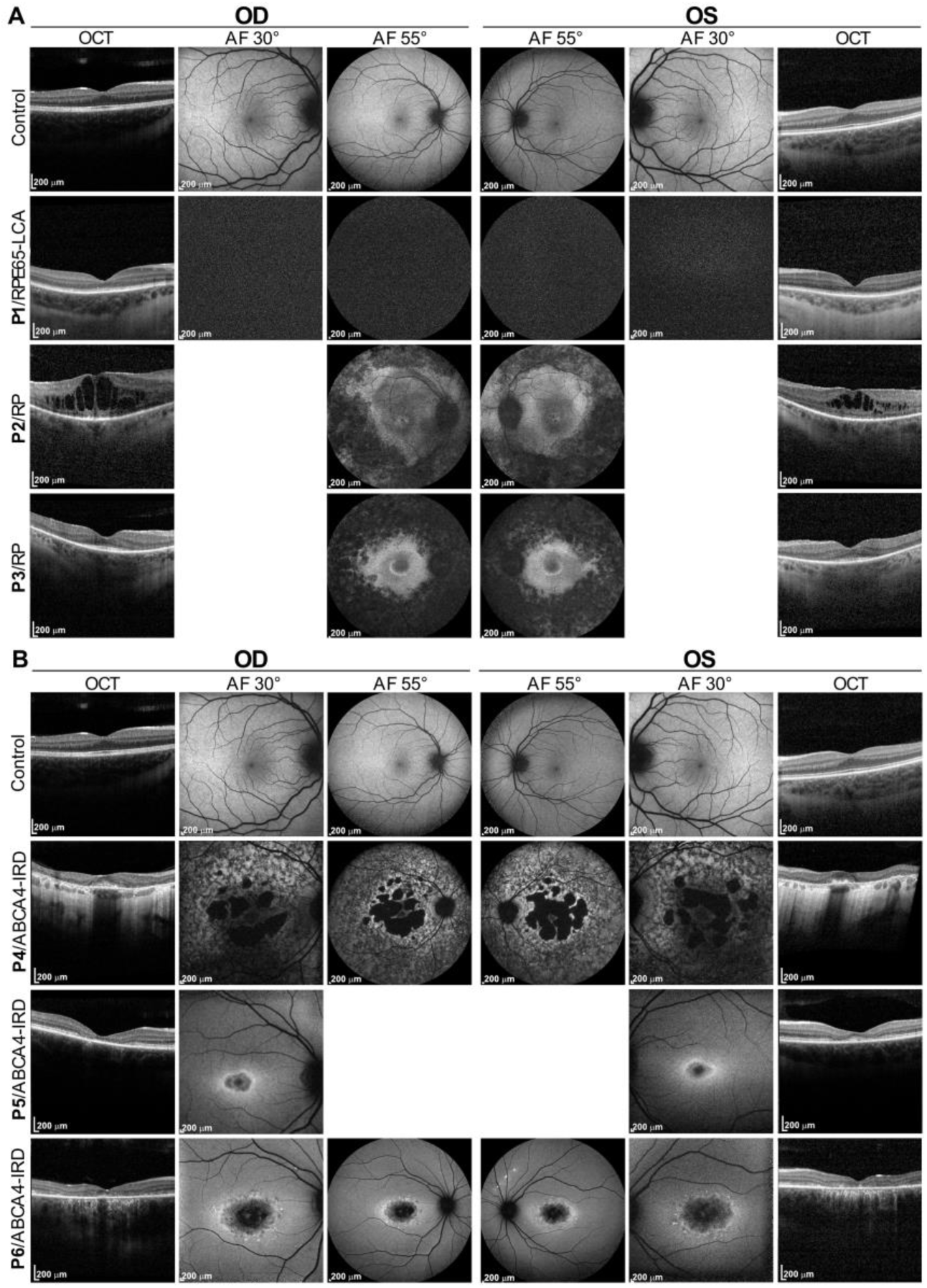
Clinical phenotype of patients. **A**, Top panel shows optical coherence tomography (OCT), 30° and 55° fundus autofluorescence (AF) images of a healthy subject (control). Three lower panels show the corresponding images of three individuals (P1-P3) with confirmed *RPE65*-associated retinal disease (RPE65-LCA and RP). P1 shows no autofluorescence due to severe *RPE65* deficiency. **B**, OCT and AF images of one healthy subject (control) and three individuals (P4-P6) with confirmed *ABCA4*-associated retinal disease (STGD1). AF, autofluorescence. OD, oculus dexter. OS, oculus sinister.

**Table 1:** Genetic diagnosis for each patient and sequencing methods.

| Patient | Gene and mutation | Comments |
| --- | --- | --- |
| P1 | *RPE65*, c.11+5G>A, heterozygous;<br>*RPE65*, c.726-2A>T, heterozygous | Exome sequencing<sup>A</sup> (Twist Human Core Exome + Twist Human RefSeq panel); Compound heterozygosity confirmed by segregation analysis<sup>B</sup> |
| P2 | *RPE65*, c.245+5A>G, heterozygous;<br>*CRB1*, c.585C>G, heterozygous;<br>*RP1L1*, c.2465G>A, heterozygous | Gene panel diagnostics comprising 42 known IRD genes; WES did not reveal additional information; Sibling of P3 |
| P3 | No genetic testing result available | Clinical phenotype of RP; Sibling of P2 |
| P4 | *ABCA4*, c.1937+1G>A, heterozygous;<br>*ABCA4*, c.5603A>T, heterozygous;<br>*ABCA4*, c.1009T>C, heterozygous | Direct sequencing of the *ABCA4* gene and segregation analysis<sup>C</sup> |
| P5 | *ABCA4*, c.2966T>C, heterozygous;<br>*ABCA4*, c.5603A>T, homozygous | WES; Sibling of P6 |
| P6 | *ABCA4*, c.2966T>C, heterozygous;<br>*ABCA4*, c.5603A>T, homozygous | WES; Sibling of P5 |

<sup>A</sup>Twist Human Core Exome combined with Twist Human RefSeq panel; <sup>B</sup>c.11+5G>A was detected in the mother and c.726-2A>T was detected in the father of P1; <sup>C</sup>Mother of P4 is carrier of the heterozygous missense mutations c.5603A>T and c.1009T>C. The father of P4 was not available for segregation analysis.

All patients, except for P3, had previously undergone molecular genetic testing of whom only P1 and P4 had a definite molecular genetic diagnosis (compound heterozygosity), which was confirmed by segregation analysis.

P1 is a 6-year-old male who has been severely visually impaired from birth and received a genetic diagnosis of *RPE65*-associated LCA at the age of 5. He was treated with voretigene neparvovec at the age of 6 (15). P2 and P3 are two female siblings who first experienced impaired vision (night blindness and in the further course of the disease visual field restriction) in their early 20’s with the clinical diagnosis of rod-cone dystrophy (RP) lacking a definite genetic diagnosis. In 2013, an initial molecular genetic testing via gene panel sequencing in P2 comprising 42 known IRD genes revealed one heterozygous variant of unknown significance in the *RPE65*, the *CRB1* and the *RP1L1* gene, respectively (Tab. 1). WES performed in 2021 did not provide any additional information.

P4 is a 49-year-old female who first experienced visual deterioration at the age of 33 representing a later onset form of STGD with pronounced macular atrophy but foveal sparing (Fig. 4). P5 is a 28-year-old female and P6 is her 31-year-old brother, both diagnosed with macular dystrophy at the ages of 19 and 21, respectively.

Potentially pathogenic compound heterozygous mutations in *ABCA4* and *RPE65* were found in all STGD1 patients (P4-P6) and in the P1 patient, respectively (Tab. 1).

We performed the CATALYTEC on PBMCs of all these patients and analyzed the isolated RNA by RT-PCR. P1 carries c.11+5G>A within the splice donor site of exon 1 in the *RPE65* gene. According to RT-PCR analysis, this mutation does not severely affect mRNA splicing at that position (Fig. 5B). For the second mutation affecting the canonical consensus sequence of the splice acceptor site in intron 7 (c.726-2A>T, Fig.5A), RT-PCR analysis revealed a clear difference in the band pattern compared to the unaffected control sample, experimentally confirming the splicing divergence. Sanger sequencing of these bands revealed partial skipping of exon 8 and inclusion of intron 7 (Fig. 5B, C).

**Figure 5.**
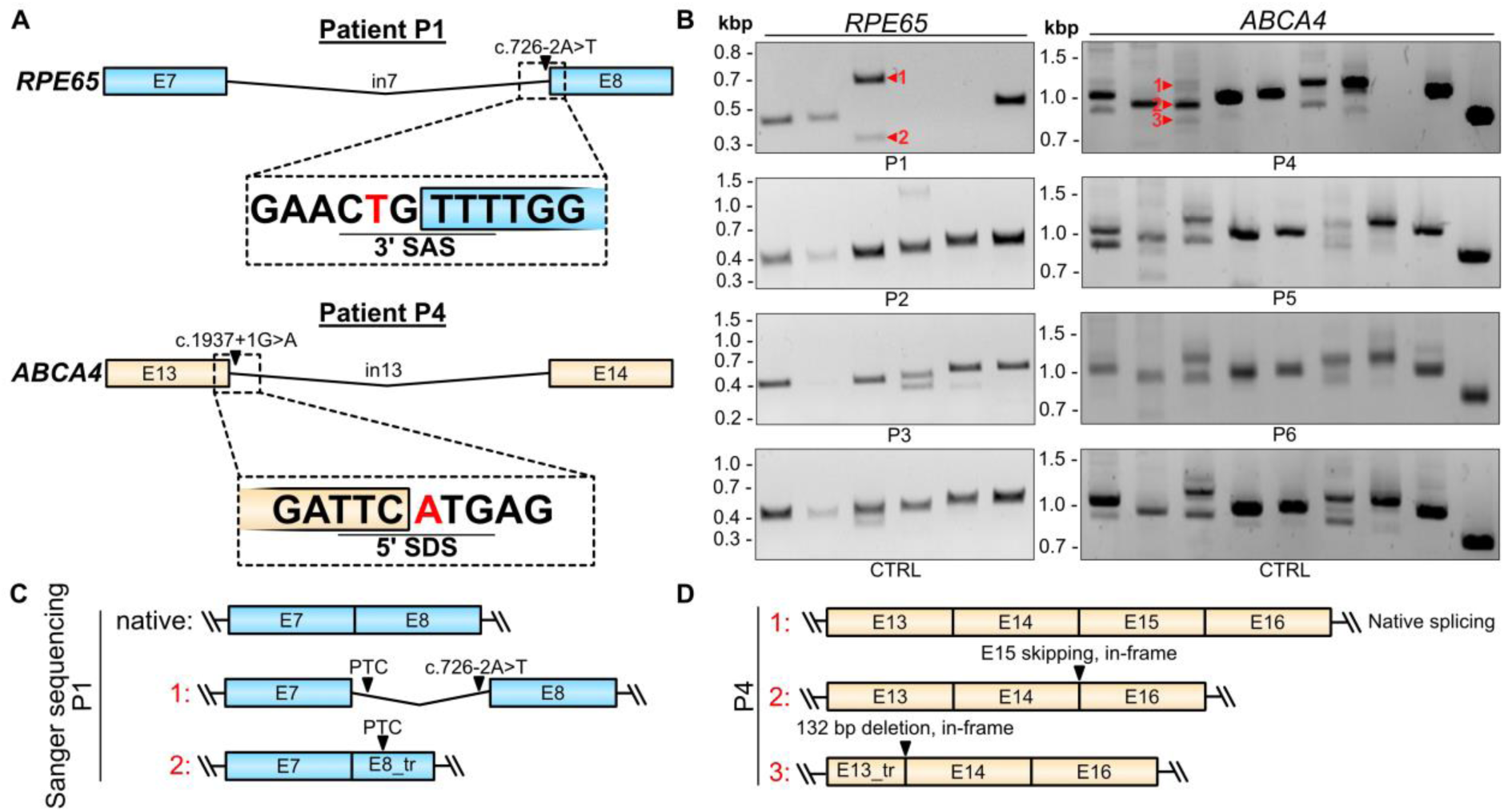
RPE65 and ABCA4 transcript analysis in patient samples. **A**, Scheme highlighting the position of the c.726-2A>T mutation in *RPE65* (patient P1; blue) and c.1937+1G>A mutation in *ABCA4* (patient P4; orange). Colored boxes represent exons, the line in between represents the intron. **B**, RT-PCR results for patients P1 – P6 in comparison to a healthy human control sample (CTRL). **C**, Scheme summarizing Sanger sequencing result of the numbered bands 1 and 2 of sample P1 shown in B. PTC, premature termination codon. **D**, Scheme summarizing the results of RT-PCR analysis of the numbered bands 1 – 3 from sample P4, as indicated in B.

The c.1937+1G>A mutation of P4 disrupts the splice donor site of exon 13 in *ABCA4* (Fig. 5A). A previous case study has associated this mutation with severe phenotypes, including STGD and macular degeneration (16). A recent publication reported that it causes aberrant splicing by activating cryptic splice donor sites within exon 13 and intron 13, ultimately resulting in a 132 bp in-frame deletion (17). This conclusion was based on data obtained with a minigene assay. Potential consequences at the protein level, such as p.[(Y.603_S646del, F647*)], have not yet been experimentally validated. For this mutation, we visually detected an increased ratio of alternative exon 15 splicing in comparison to healthy control samples in RT-PCR (Fig. 5B, D). As described in Table 1, this alternatively spliced *ABCA4* variant leads to an in-frame skipping of exon i) 15. However, the significance of this deletion for ABCA4 protein expression, stability and function remains unclear and is outside the focus of this study. For all other patients carrying potentially pathogenic mutations in IRD genes analyzed herein, no obvious effects on splicing could be detected (Fig. 5B).

Taken together, these results suggest that our CATALYTEC is in principle suitable for the detection and quantification of splicing mutations in patients’ PMBCs.

## Discussion

Here, we established CATALYTEC, a robust and simple approach for CRISPRa-based activation of genes in human PBMCs and skin fibroblasts suitable for diagnostic purposes.

CATALYTEC fulfills various criteria for use in diagnostics:

i. Due to its simplicity, it is easy to implement in routine diagnostics.
ii. It can be utilized to detect novel mutations in known genes, which might have been overlooked in the past due to technical limitations.
iii. It can be used to validate the proposed pathogenicity of detected mutations.
iv. It is without further modifications transferable to other genetic disorders where biopsy of the affected tissue from the patients is not possible.

In recent studies, similar approaches were developed to activate the *MPZ* and *SPAST* genes associated with Charcot-Marie-Tooth disease and hereditary spastic paraplegia, as well as the *CRB1* gene associated with retinitis pigmentosa, in human skin fibroblasts (18, 19). Compared to these studies, our method offers additional key advancements. We demonstrate for the first time that dCas9-VPR can effectively activate disease-associated genes in PBMCs readily isolated from patient blood samples. Skin fibroblasts have the disadvantage that their isolation requires invasive skin punching, which is more elaborate and associated with reduced patient compliance. Therefore, genetic testing of human skin fibroblasts is less suitable for broad routine diagnostics compared to PBMCs. Additionally, the isolation and cultivation of skin fibroblasts is more time-consuming. With our CATALYTEC approach, we achieved a turnaround time from blood collection to first RT-PCR results of 48-72h, ultimately increasing the convenience for patients and the healthcare system.

We have applied CATALYTEC to several common genes associated with IRDs, a heterogeneous group of genetic diseases affecting the retina. In particular, we have proven the ability of our approach to activate large genes like *ABCA4* and *MYO7A*. This is of great importance as deep intronic mutations in large genes are not covered by WES or, when identified by WGS, cannot be correctly interpreted in terms of their potential impact on mRNA splicing.

Our data shows that the level of transcriptional activation obtained with CATALYTEC is high enough to be combined with the most commonly used readout methods (RT-PCR, RT-qPCR, short- and long-read RNA sequencing). RT-PCR analysis provided the first evidence that CATALYTEC can be used to detect pathogenic splice mutations in PBMCs of IRD patients. Additionally, we show that short- and long-read sequencing can be used to analyze the expression and alternative splicing of transcriptionally activated genes in PBMCs. However, a meaningful and significant result for diagnostic purposes necessitates a more detailed analysis of isoforms, which could be achieved through higher coverage of the respective target genes.

We demonstrate that the splicing patterns of activated IRD genes in human PBMCs and fibroblasts are very similar to those observed in human retinas and retinal organoids. Yet, we cannot exclude the possibility that other activated genes in PBMCs might have different splicing patterns than in the cells in which these genes are naturally expressed. In addition, rare tissue-specific splicing events may have been missed by RT-PCR. More detailed investigations using short- and long-read sequencing of the human retina and organoid transcriptomes could provide valuable insights in this regard.

An attractive alternative to PBMCs would be an application of the CATALYTEC protocol in buccal epithelial cells, as their collection is technically even simpler and less invasive than blood sampling. It remains to be seen whether this goal can be achieved using CRISPRa or other methods for introducing DNA or RNA into these cells.

In summary, we provide proof of concept for a CRISPRa-based approach to activate IRD-associated genes in PBMCs in sufficient quantities. The resulting transcripts can be detected and quantified using standard methods and analyzed for structural variants and splicing defects. The CATALYTEC method is universally applicable, can be easily adapted to other target genes, and could help closing important gaps in the diagnosis of inherited (retinal) diseases.

## Methods

### Sex as a biological variable

Our study examined men and women, as gender should not be an exclusion criterion due to the diagnostic background of the study.

### Plasmids

The pcDNA3.1-CMV-dCas9-VPR and plasmids for LV production (pMDL, pRSVRev and pMD2.G) were obtained from Addgene (#63798, #12251, #12253, #12259). Expression of dCas9-VPR in pcDNA3.1 vectors was driven by the cytomegalovirus (CMV) promoter. In lentiviral CRISPRa systems, the dCas9-VPR expression was induced by either CMV, spleen focus-forming virus (SFFV) or elongation factor 1α (EF1α) promoter.

Single-guide RNA cassettes were synthesized (Azenta, IDT) and inserted using standard cloning techniques. Sequences are shown in Suppl. Table 3. All plasmids were sequenced before use (Eurofins Genomics/Microsynth).

### Cell Culture

HEK293T cells (Takara Bio) were maintained in DMEM (high glucose, Thermo Fisher Scientific) supplemented with 10% FBS (Superior, Sigma-Aldrich) and 1% penicillin/streptomycin (P/S, Thermo Fisher Scientific) at 37 °C, 10 % CO_2_.

Human skin fibroblasts (adult, Sigma-Aldrich) were cultured in DMEM (low glucose) supplemented with 10% FBS and 1% P/S at 37 °C, 5% CO_2_.

PBMCs were cultured in RPMI-1640 (Thermo Fisher Scientific) supplemented with 10% FBS at 37 °C, 5% CO_2_.

### Collection and stimulation of PBMCs

PBMCs were isolated through density centrifugation of the collected whole blood. The isolated buffy coat was washed with PBS and resuspended in RPMI-1640, supplemented with 10% FBS and 1X phytohemagluttinin-L (PHA-L, Thermo Fisher Scientific). The cells were incubated at 37 °C and 5% CO_2_ for 20 h before being used for nucleofection.

### Isolation of human retinas

Human donor eyes without cornea and lens were received from the eye bank of the University Hospital Zurich in ice-cold PBS. Retina and eyecups were separately flat mounted after making four incisions. Retina samples were collected from the nasal periphery and the macular region. The isolated retina was snap-frozen and stored at -80 °C until further processing.

### Transfection of cell lines and primary cells

HEK293T were transfected with either Xfect^TM^ (Takara Bio) or Lipofectamine 3000 (Invitrogen) according to the manufacturers’ instructions and harvested 48 h after transfection.

PHA-L-stimulated PBMCs (2.0-2.5x10^^7^ cells per reaction) were nucleofected with the P3 Primary Cell 4D-Nucleofector Kit (Lonza). After application of program EO-115, 500 µL of pre-equilibrated culture medium was added to the cuvette and the suspension was immediately transferred to a pre-equilibrated 12-well plate. Nucleofected PBMCs were incubated for 24 h at 37 °C, 5% CO_2_.

For nucleofection of fibroblasts (2x10^^6^ cells per reaction) the P2 Primary Cell 4D-Nucleofector Kit (Lonza) was used. After application of program CZ-167, 500 µL of pre-equilibrated culture medium was added to the cuvette and the suspension was immediately transferred to a pre-equilibrated 6-well plate. Nucleofected fibroblasts were incubated for 24 h at 37 °C, 5% CO_2_.

### RNA isolation

HEK293T and fibroblasts were washed with PBS and lysed with RLT plus buffer (Qiagen) supplemented with 10 µL/mL β-mercaptoethanol (Carl Roth). The cell lysate was transferred into safe-lock tubes (Eppendorf) and homogenized with a mixer mill (Retsch) at 30 Hz for 1 min. After centrifugation, RNA was isolated according to the manufacturer’s protocol of the RNeasy Plus Mini Kit (Qiagen).

For PBMCs, the cell were pelleted and lysed with RLT plus buffer supplemented with 10 μL/mL β-mercaptoethanol. Subsequently, the lysate was added to QIAshredder homogenization columns (Qiagen) and centrifuged according to the manufacturers’ protocol. After homogenization, the RNeasy plus Mini Kit was used to isolate the RNA.

Snap-frozen retinas were lysed in buffer RLT supplemented with 10 µL/mL β-mercaptoethanol. The retinas were homogenized through a 21G needle. Afterwards, the RNA was isolated with the RNeasy Plus Mini Kit.

RNA isolation from human retinal organoids was done with the Direct-zol DNA/RNA miniprep kit (Zymo) according to the manufacturer’s instructions.

To avoid gDNA contamination, an on-column DNase I digest (Qiagen) was performed for every RNA isolation.

### Two-step RT-PCR

First-strand cDNA for two-step RT-PCR analysis of HEK293T, PBMCs and fibroblasts was produced with Maxima H Minus Reverse Transcriptase (Thermo Fisher Scientific) according to the manufacturers’ protocol. First-strand cDNA from retinal RNA was synthesized with the M-MLV reverse transcriptase (Promega).

Subsequent second-strand synthesis and PCR amplification was performed with the Q5 Hot Start High Fidelity Polymerase (New England Biolabs). Primers used for amplification are listed in Suppl. Table 4. Results were visually analyzed through agarose gel electrophoresis (1% w/v). Amplified bands were isolated with the QIAquick Gel Extraction Kit (Qiagen) and sent for Sanger sequencing (Microsynth AG, Eurofins).

### RT-qPCR

For quantitative real-timer PCR, RNA was reverse transcribed into cDNA with the RevertAid First Strand cDNA synthesis Kit (Thermo Fisher Scientific). The SYBR^TM^ Green PCR Master Mix (Thermo Fisher Scientific) was used to prepare the samples according to the manufacturers’ instructions. For amplification and analysis, the MicroAmp^TM^ Fast Optical 96-Well Reaction Plate and QuantStudio 3 RT-PCR system and software (Thermo Fisher Scientific) were used. Expression levels were normalized to *ALAS1*. Primers used for RT-qPCR analysis are listed in Suppl. Table 5.

### Optical coherence tomography (OCT) and autofluorescence (FAF) imaging

Retinal cross sections were obtained with spectral-domain optical coherence (SD-OCT) tomography using the Heidelberg Spectralis® OCT (Heidelberg Engineering). FAF (488 nm) was obtained using the same device.

### Lentivirus production

For production of lentiviral vectors, HEK293T were transfected with pMDL, pRSVRev, pMD2.G as well as the transgene plasmid plasmids via calcium phosphate method. The transfected cells were incubated for 48 h at 37 °C, 10% CO_2_. Subsequently, the cell culture medium was collected and filtered with a 0.45 µm filter unit (VWR). The cells were supplied with fresh culture medium and incubated for additional 24 h. The filtered medium was centrifuged at 19400 rpm and 17 °C for 2 h (Beckman Coulter). The pellet was suspended in 250 µL HBSS (Thermo Fisher Scientific) and stored at 4 °C overnight. The described procedure was repeated with the cell culture media, added the day before. The first and second harvest were combined and concentrated with a sucrose cushion centrifugation at 21000 rpm, 17 °C for 2 h. The resulting pellet was suspended in 70 µL HBSS and mixed for 45 min at 1400 rpm. Final aliquots of 5 µL were stored at -80 °C.

Titer determination was performed with a RT-qPCR Lentivirus Titer Kit (Applied Biological Materials Inc.) according to the manufacturers’ instructions.

### Generation of human retinal organoids

Human retinal organoids were differentiated from the human derived iPSCs (F49B7). Pluripotency markers and germ layer differentiation potential was determined as previously described (20). IPSCs were seeded in matrigel-coated 6-well plates (Corning) and cultured in mTeSR plus medium (STEMCELL) at 37 °C, 5% CO_2_. The iPSCs were passaged using 0.5 mM EDTA (pH 8.0, Thermo Fisher Scientific).

Differentiation of iPSCs into human retinal organoids was performed according to the protocol developed by Kim et al (21), with some modifications. All organoids at the different maturation stages were cultured in a humidified incubator at 37 °C, 5% CO_2_.

On day 0, the iPSCs were dissociated using 0.5 mM EDTA. The aggregates were suspended in cold Matrigel (GFR, Corning) and incubated at 37 °C for 20 min. Afterwards, the iPSC/Matrigel aggregates were dispersed in neural induction medium (DMEM/F12 with neurobasal medium (1:1) supplemented with 1% B27 (incl. vitamin A supplement), 0.5% N-2 supplement, 0.1 mM β-mercaptoethanol, 2 mM GlutaMax and 1% P/S (all Thermo Fisher Scientific)) and cultivated in ultra-low adherent 6-well plates (Costar®, Corning). On day 5, floating cysts were seeded in matrigel-coated 6-well plates. On day 15, cysts were detached by adding dispase (0.5 mg/mL in DMEM/F12, STEMCELL), washed with DMEM/F12 and further cultured in retinal differentiation medium (DMEM/GlutaMax supplemented with F12 nutrition mix (3:1), 2% B27 (without vitamin A), 1% non-essential amino acids (NEAA) and 1% P/S). On day 25, the immature retinal organoids were transferred to retinal maturation medium (DMEM/GlutaMax supplemented with F12 nutrition mix (3:1), 8% FBS, 2% B27 (without vitamin A), 1% NEAA, 1% antibiotic/antimycotic and 1% 100 mM taurine (Sigma-Aldrich).

On day 230, mature organoids were used for RNA isolation, AAV transduction and subsequent experiments.

### Short-read RNA sequencing of healthy blood samples

For short read mRNA sequencing, 100 ng of DNase treated total RNA (RIN > 8) was processed with the TruSeq Stranded mRNA Prep Kit (Illumina) including poly(A) selection. Indexes were added with the Illumina RNA UDI 384 v2 kit (IDT). 150 bp paired end sequencing was performed with a NovaSeq X system (Illumina). Sequenced reads were aligned to the human reference genome (GRCh38) and were counted on gene-level with Rsubread (version 2.18). Differentially expressed genes (DEGs) were determined between control and treated samples employing edgeR (v4.2.1) and limma (v3.60.3). Gene set enrichment analysis of DEGs was carried out with fgsea (v1.30.0). Sashimi blots were generated with Integrative Genomics Viewer (IGV, version 2.17.4 03/26/2024).

### Long read RNA sequencing of healthy blood samples

For PacBio long-read sequencing healthy blood samples were sent to an external sequencing facility (Bioscientia). Sequencing was performed on a PacBio Revio device (Pacific Biosciences). The Isoseq (v4.0.0; https://isoseq.how/) workflow was utilized to analyze Hifi reads and quantify gene expression, as well as, extract different transcript isoforms. Mapped long reads were visualized in IGV.

### WES of patient-derived blood samples

Genomic DNA, isolated from collected blood samples, was fragmented, and the coding exons of the analyzed genes as well as the corresponding exon-intron boundaries were enriched using Roche/KAPA sequence capture technology (KAPA HyperExome Library) and sequenced using an Illumina NovaSeq 6000 system. The requested gene panel was extracted from the WES data. The target regions were sequenced with an average coverage of 337x. For more than 99% of the target regions a 15-fold coverage was obtained. Putatively pathogenic differences between the wildtype sequence (human reference genome according to UCSC Genome Browser: hg19, GRCh37) and the interpreted patient’s sequence were assessed using an internally established quality system. Variants, which did not pass the quality threshold, were verified using conventional Sanger sequencing. Variants listed as additional, putatively relevant variants were not routinely validated. Identified variants were compared to literature and external as well as internal allele frequency databases (e.g. gnomAD). In addition, *in silico* analysis of the identified variants with regard to functional relevance, conservation and splice effects was performed using bioinformatic prediction programs (e.g. SpliceAI, MaxEntScan). The variants were classified using the current ACMG guidelines (22).

### Long-read WGS of patient-derived blood samples

Genomic DNA, isolated from collected blood samples, was fragmented, and a PCR-free library was prepared using SMRTbell prep kit 3.0 (PacBio). Long-read whole-genome sequencing was done using a PacBio Revio system at an average coverage of approx. 30-fold. Putatively pathogenic differences between the wildtype sequence (human reference genome according to UCSC Genome Browser: hg19, GRCh37) and the patient’s sequence mentioned and interpreted in this report were assessed using an internally established quality system.

Identified variants were compared to literature and external as well as internal allele frequency databases (e.g. gnomAD). In addition, *in silico* analysis of the identified variants with regard to functional relevance, conservation and splice effects was performed using bioinformatic prediction programs. The variants are classified using the current ACMG guidelines (22).

The genome data were filtered with respect to autosomal recessive, autosomal dominant and X-linked mode of inheritance for very rare potentially pathogenic homozygous/putative compound heterozygous/heterozygous and hemizygous changes. For the filtered variants, a literature-based comparison using human mutation databases (e.g. HGMD, ClinVar) was performed according to the provided clinical information of the patient. In addition, the data were compared to public and internal allele frequency databases. Furthermore, the *in-silico* scores of bioinformatic prediction programs were also taken into account. The NGS data were not analyzed for potentially pathogenic variants in genes not related to the requested indication. Additional, putatively relevant variants and carriership findings are not reported routinely.

### Statistical analysis and reproducibility

All values are given as mean ± SEM. Statistics were performed with GraphPad PRISM (GraphPad Software, v10.0.2).

### Study approval

The patients (P1-P6) involved in this project presented at the Department of Ophthalmology at LMU Munich and research including patient samples were approved by the local ethics committee (ethics vote nr. 19-0226). Clinical research and publication of clinical imaging data was approved by the local ethics committee (ethics vote nr. 22-0897). Written informed consent for the use of patient samples and clinical imaging was received prior to participation and has been retained.

Human eyes, used for human retina analysis, were donated and collected in collaboration with the Eye Clinic Zurich and were approved by the local ethics committee (BASEC-Nr: PB_2017-00550 and 2020-01856).

All procedures with human samples and donations adhered to the tenets of the Declaration of Helsinki.

## Data availability

Values for all data points shown in graphs are reported in the Supporting Data Value file.

Sequencing data of human samples have not been deposited in a public repository because this could compromise the privacy of the research participants, but they are available from the corresponding author upon request.

## Author contributions

V.J.W., K.S.H., S.M. and E.B. designed the research studies. V.J.W., M.J.G., A.R., K.S.H., H.J.B. and D.Y.O. conducted the experiments and acquired data. T.H., V.J.W. analyzed data. F.B. and I.M. provided human retina samples. V.J.W., M.J.G and E.B. wrote the manuscript. E.B., S.M. and M.B. acquired funding. S.M. and M.J.G. supervised experiments with human patient samples. E.B. supervised the project. All authors contributed to the final manuscript.

## Acknowledgements

This research study was funded by the Helmut-Ecker Stiftung (to E.B.), Novartis Stiftung (to S.M.), Iten-Kohaut Stiftung (to E.B.) and the Swiss National Science Foundation (to E.B., 320030E_221942).

We thank Bioscientia Healthcare GmbH for performing the library preparation and the sequencing for the PacBio long-read RNA sequencing approaches.

We thank Claudia Matter for the processing of human eye donations.

Library preparation and sequencing for short-read RNA sequencing was performed by the Functional Genomics Center Zurich (FGCZ).

Figure 3A and the graphical abstract were created with BioRender.com (https://app.biorender.com/).

