## Supplementary figures and tables for "CRISPRa-mediated activation of genes associated with inherited retinal dystrophies in acutely isolated human cells for diagnostic purposes"

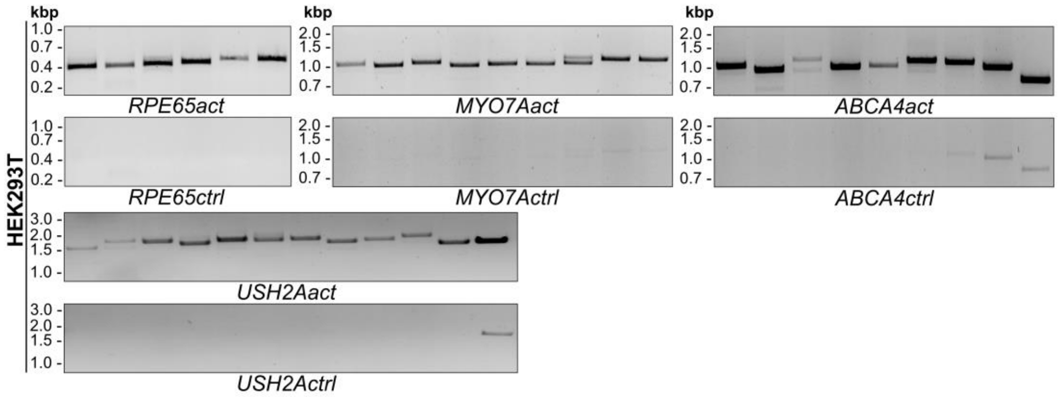


**Suppl. Fig. 1 Extended RT-PCR analyses of HEK293T containing control samples without transcriptional activation (related to Fig. 1).** For further details, see main text and Fig. 1.

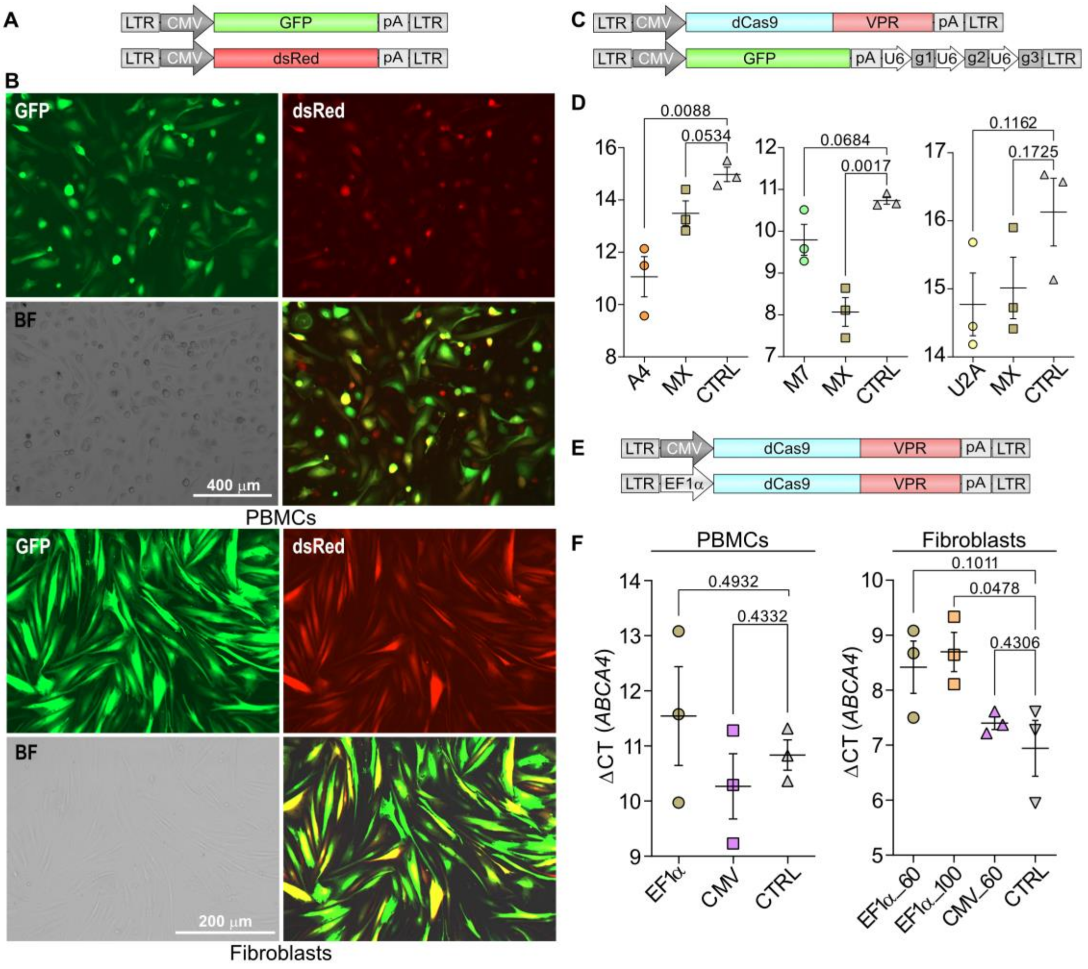


**Suppl. Fig. 2 Lentiviral transduction and optimization of CRISPRa in human PBMCs and fibroblasts. A,** Scheme depicting the LV vector expression cassettes used for evaluation of co-transduction. GFP, green fluorescent protein. LTR, long terminal repeats. **B,** Microscopic imaging of human PBMCs and fibroblasts co-transduced with LVs expressing either GFP or dsRed. For this experiment, PBMCs have been cultured for two and fibroblasts for one week before transduction. BF, bright field. **C,** Scheme showing lentiviral expression plasmids for dCas9-VPR mediated gene activation. **D,** RT-qPCR showing target gene expression after dual transduction of HEK293T. **E,** Scheme depicting LV expression cassettes for promoter evaluation. EF1α, Elongation factor 1α promoter. **F,** RT-qPCR showing *ABCA4* expression after transduction of PBMCs with LVs expressing dCas9-VPR under control of CMV or EF1α promoter (left) and with different multiplicities of infections (right). Values are given as mean ± SEM. Statistics were calculated with Student's t-test.

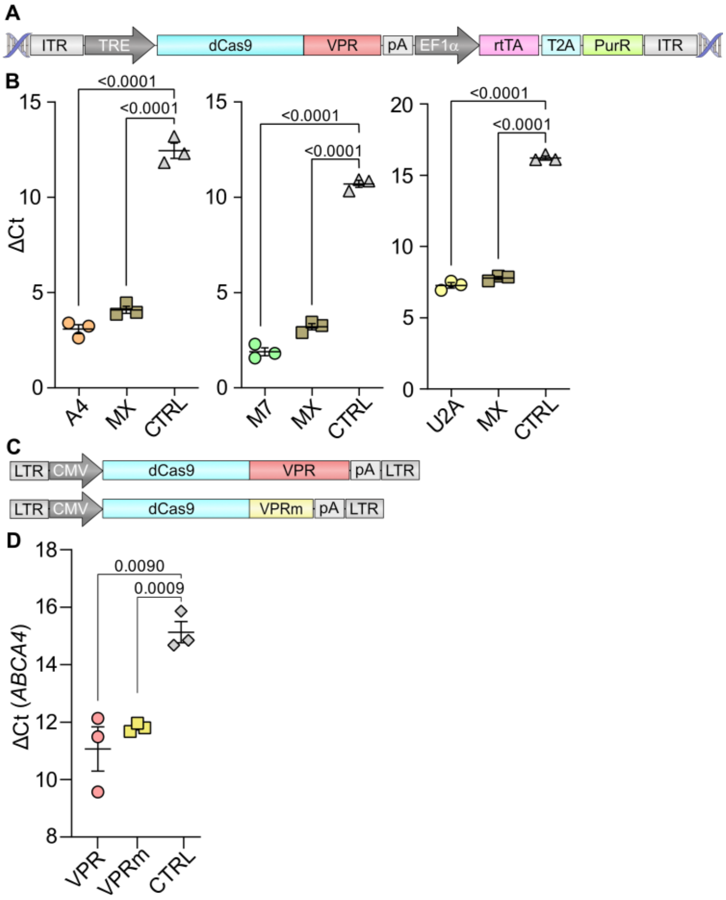


**Suppl. Fig. 3 Generation of HEK293T cells stably expressing dCas9-VPR and evaluation of a smaller CRISPRa module. A,** PiggyBac gene expression vector used for integration in the HEK293T genome. ITR, inverted terminal repeats; TRE, tetracycline responsive element; rtTA, reverse tetracycline-controlled transactivator. **B,** RT-qPCR result for each gene activation. A4, *ABCA4*; M7, *MYO7A*; U2A, *USH2A*; MX, multiplexing with all three sgRNA cassettes. **C,** Plasmid scheme for LVs expressing dCas9 in combination with VPR or VPRmini (VPRm). **D,** RT-qPCR result for *ABCA4* expression after co-transduction of HEK293T. All values are shown as mean ± SEM. Statistics were calculated with Student's t-test.

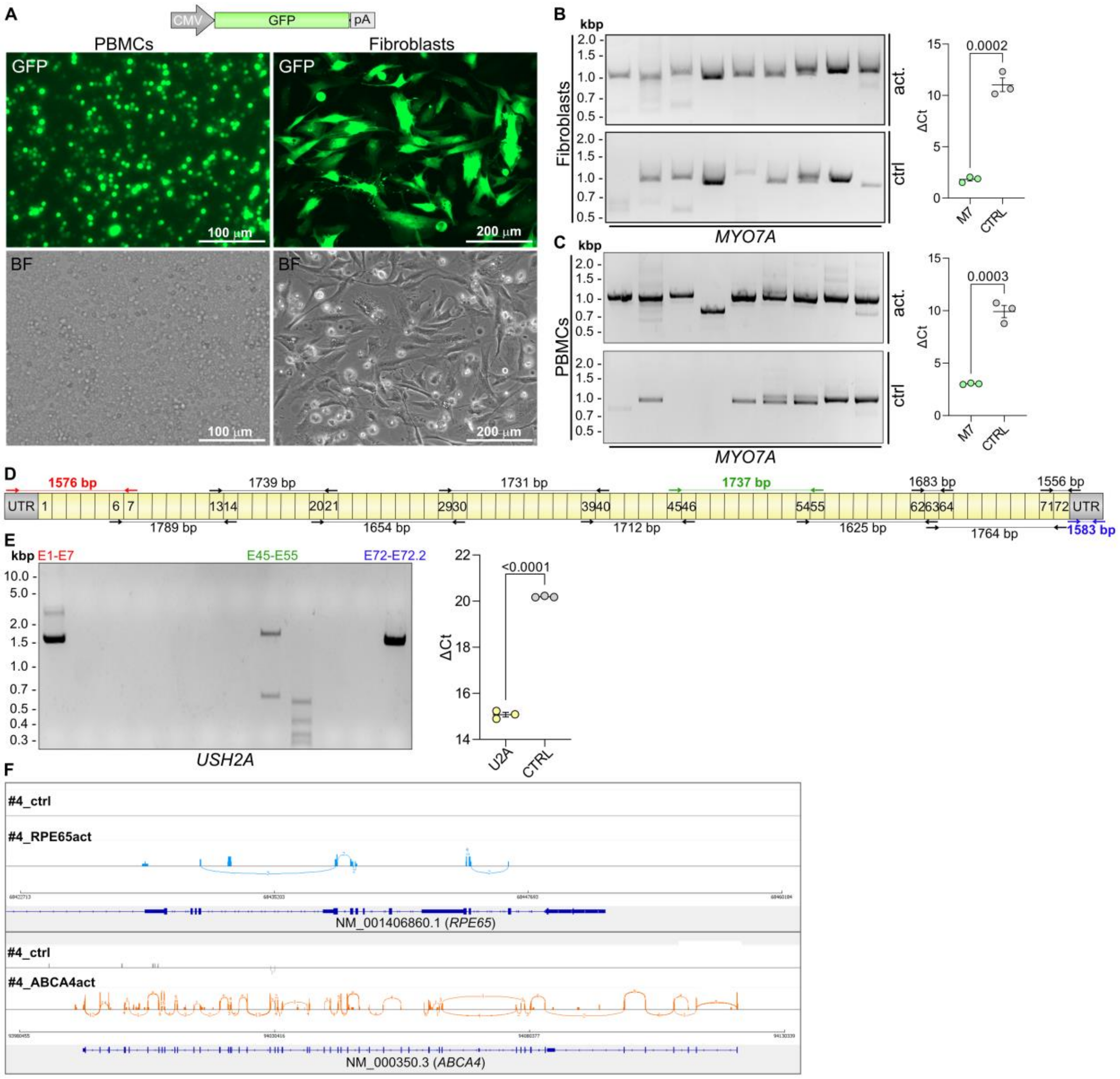


**Suppl. Fig. 4 Nucleofection of primary target cells and evaluation of transcriptionally activated *MYO7A* and *USH2A*. A,** Upper panel: GFP reporter plasmid used for nucleofection. Lower panel: Live imaging of PBMCs and fibroblasts after nucleofection with the GFP reporter plasmid. **B,** RT-PCR (left) and RT-qPCR (right) for *MYO7A* in fibroblasts after nucleofection and transcriptional activation. **C,** RT-PCR (left) and RT-qPCR (right) for *MYO7A* in PBMCs after nucleofection and transcriptional activation. **D,** Primer design for the RT-PCR analysis of the *USH2A* transcript. **E,** *USH2A* amplifications (left) of exon 1-7 (E1-E7, red), exon 45-55 (E45-E55, green) and exon 72-72-2 (E72-E72.2, blue) as indicated in D. RT-qPCR (right) for *USH2A* after nucleofection of PBMCs with CRISPRa plasmids. All values are shown as mean ± SEM. Statistics are calculated with Student's t-test. **F,** Sashimi blot for transcriptionally activated *RPE65* (blue) and *ABCA4* (orange) compared to untreated control PBMCs (grey). Blue and orange bars represent the respective exon coverage. The numbered curved lines show the number of reads spanning of exon-exon junctions. Dark blue schemes highlight the main reference transcript of the respective genes.

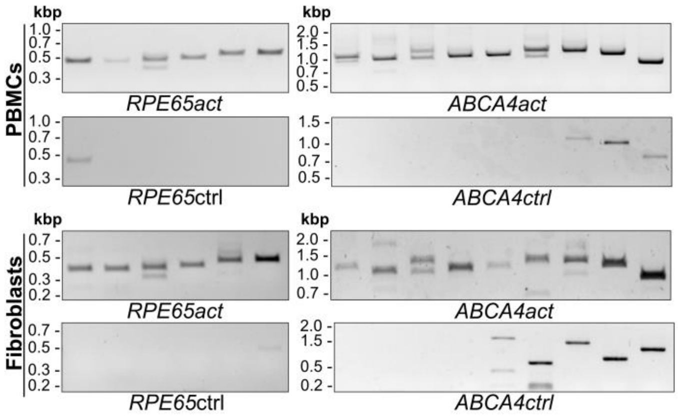


**Suppl. Fig. 5 Extended RT-PCR analyses of PBMCs and fibroblasts containing control samples without transcriptional activation (related to Fig. 2)**. For further details, see main text and Fig. 2.

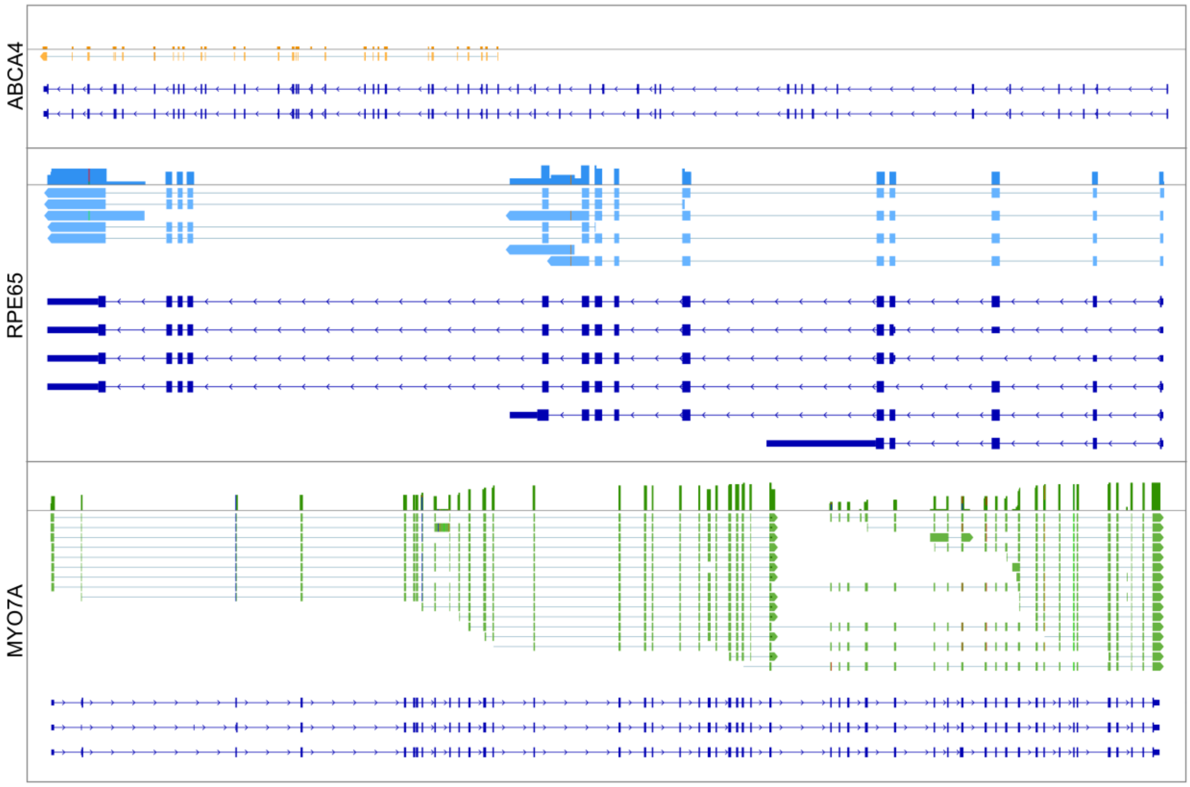


**Suppl. Fig. 6 Consensus sequences for *ABCA4*, *RPE65* and *MYO7A* after long-read sequencing.** Consensus sequences for *ABCA4* (orange), *RPE65* (blue) and *MYO7A* (green) after transcriptional activation in human PBMCs. The colored bars on top of each consensus sequence panel represents the exon coverage. Detected mutations are labelled in red, bright green orange or blue. Dark blue schemes highlight the reference transcripts of the respective genes.

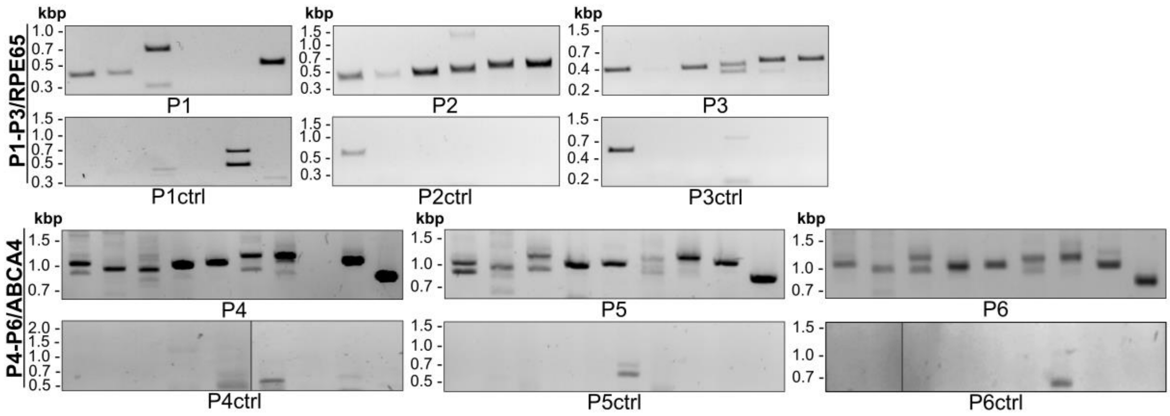


**Suppl. Fig. 7 Extended RT-PCR analyses of patient samples containing control samples without transcriptional activation (related to Fig. 5).** For further details, see main text and Fig. 5.

**Suppl. Table 1: Sanger sequencing result of *ABCA4* amplicons**

| Exon range | Band identity |
|---|---|
| E1 – 8 | Upper: Canonical sequence (Exon 1-8)<br>Lower: Exon 3 skipping (PTC) |
| E7 – 10 | Constitutive splicing |
| E11 – 16 | Upper: Canonical sequence (Exon 11-16)<br>Lower: Exon 15 skipping (in frame) |
| E15 – 21 | Constitutive splicing |
| E20 – 27 | Constitutive splicing |
| E26 – 35 | Upper: Canonical sequence (Exon 26-35)<br>Lower: Exon 28+29 skipping (PTC) |
| E34 – 42 | Constitutive splicing |
| E41 – 48 | Constitutive splicing |
| E47 – 50 | Constitutive splicing |

**Suppl. Table 2: Sanger sequencing result of *RPE65* amplicons**

| **Exon range** | **Band identity** |
|---|---|
| E1 – 4 | Constitutive splicing I |
| E3 – 6 | Constitutive splicing |
| E5 – 8 | Upper: Canonical sequence (Exon 5 - 8)<br>Lower: Exon 7 skipping (PTC) |
| E7 – 10 | Constitutive splicing |
| E9 – 13 | Constitutive splicing |
| E12 – 14 | Constitutive splicing |

**Suppl. Table 3: sgRNA sequences**

| sgRNA | Sequence |
|---|---|
| sgRPE65_1 | CCCAAATAAAGCCAAGCATC |
| sgRPE65_2 | GGGGTGACTGGGATCAGCTC |
| sgRPE65_3 | CCTCCCTCAGCTGAAGGGGT |
| sgABCA4_1 | TTACACATAAGCCCTTAGTT |
| sgABCA4_2 | TACCTAAAGACATCCCCCTC |
| sgABCA4_3 | CCCTAAAGACACAAGTGGCT |
| sgMYO7A_1 | GGCGGTGTCCTGCAAACAAC |
| sgMYO7A_2 | CAGAGCACTGGCTGCCGACA |
| sgMYO7A_3 | GGGTGCGGTGACATCAGCGT |
| sgUSH2A_1 | TCTATCAATGGGCCAAGCTT |
| sgUSH2A_2 | AGAGGGGGGATTATAGGATA |
| sgUSH2A_3 | GAGGTACCAGCGGAAAGCTT |

**Suppl. Table 4: RT-PCR primer for *RPE65*, *ABCA4*, *MYO7A* and *USH2A* transcript amplification**

| Primer | Sequence [5’ -> 3’] |
|---|---|
| RPE65_E1_fw | GCAGTTGGTGCCAGAACTC |
| RPE65_E3_fw | CACCTGTTTGATGGGCAAGC |
| RPE65_E5_fw | ACGCTTGCACAGAGACCAAC |
| RPE65_E7_fw | TACAATTCCCCTGCAGTGACC |
| RPE65_E9_fw | TGTGGATCTCTGCTGCTGGA |
| RPE65_E12_fw | ATGCGTATGGACTTGGCTTGA |
| RPE65_E4_rev | TTGCAGGGATCTGGGAAAGC |
| RPE65_E6_rev | TTTCAATGTGGGGGTGAGCA |
| RPE65_E8_rev | GTAGTTGGCTCCCCAAAGACT |
| RPE65_E10_rev | ACTTCAGGTTGGGGAGCCTT |
| RPE65_E13_rev | CTTCCAAGGCATCTGGGTGA |
| RPE65_E14_rev | TGCTTGCTCAACTCAGTGCT |
| ABCA4_E1_fw | GAGCCAGAGGCGCTCTTAAC |
| ABCA4_E7_fw | TCCCACACTCCTAGACAGCC |
| ABCA4_E11_fw | ATACCCTGGGGAACCCAAC |
| ABCA4_E15_fw | TTACAGCGACCCATTCATCCTC |
| ABCA4_E20_fw | GTCCATCCTGACGGGTCTG |
| ABCA4_E26_fw | CTGAAGGTCACGGAGGATTCTG |
| ABCA4_E34_fw | GGCCCTATCACTAGAGAGGC |
| ABCA4_E41_fw | CCACTAAGGAGCCCATTGTTGAT |
| ABCA4_E47_fw | CTGGCCATCATGGTAAAGGGC |
| ABCA4_E8_rev | CTCTGGACCACCATTCTGCAT |
| ABCA4_E12_rev | CTTCACGTGGGGTGGTAGAG |

| Primer | Sequence [5’ -> 3’] |
|---|---|
| ABCA4_E16_rev | GGACTGTTCCCGATGTTGCT |
| ABCA4_E21_rev | TCCAACATGGCTTCCATCTCC |
| ABCA4_E27_rev | GTCGGGGGTTGACGTTTTC |
| ABCA4_E35_rev | CCGTAAGATGGCGTTGTGGG |
| ABCA4_E42_rev | GCTGGAGGTGCCTGGATAAA |
| ABCA4_E48_rev | CAGTGTGGTCTGTGTGACTGA |
| ABCA4_E50_rev | TCCATGAAAATCACACACAACGCA |
| MYO7A_E1_fw | CTGGACAGCTGCTCTGGG |
| MYO7A_E6_fw | GGGAAGACGGAGAGCACAAA |
| MYO7A_E12_fw | GTGGACAAGATCAACGCAGC |
| MYO7A_E18_fw | CGATGACTGGCAGATAGGCAA |
| MYO7A_E23_fw | GAGCTGAAGGAGAAGGAGGC |
| MYO7A_E28_fw | TTCTCGTGTCTCTCTGCGTG |
| MYO7A_E33_fw | CCAGAGGAGAACTGATGCCC |
| MYO7A_E38_fw | ACCCAAGCACACGCTGAG |
| MYO7A_E45_fw | GTATCTCCGAGGCTACCACAAG |
| MYO7A_E7_rev | ACGTGACTTTTCCAGCAGGT |
| MYO7A_E13_rev | TCGATGTGCAGCCAGTCAAT |
| MYO7A_E19_rev | ATGACTCTGTCGGTGATGGC |
| MYO7A_E24_rev | TCACCCTCGTCGTCATGGTA |
| MYO7A_E29_rev | ACAAAGGTCCTTCTCAGGCG |
| MYO7A_E34_rev | GGCCATGATCTCTGGGAAGG |
| MYO7A_E39_rev | CTGCACATATGCCTCGTCCT |
| MYO7A_E46_rev | CGTGCTTGTTGAAGTAGGCG |
| MYO7A_E49_rev | GATCAGGGTAGACTTGAGCAGG |
| USH2A_E1_fw | TGTTTGCTCTGCAGAATACTT |

| Primer | Sequence [5’ -> 3’] |
|---|---|
| USH2A_E6_fw | GTTGGTACTTCATGGGTTTCA |
| USH2A_E13_fw | GATGTGTGAGTGTGATTCCTT |
| USH2A_E20_fw | CCCAAGAACTATCTTACACTG |
| USH2A_E29_fw | AGTCGAGGACGTACAACAG |
| USH2A_E39_fw | GATGAGGTCTGGCGACT |
| USH2A_E45_fw | CAGGACTTCATGCAACCA |
| USH2A_E54_fw | TATCAGCTGAAAGCTTGCACG |
| USH2A_E62_fw | GTAGAACAGAAAGAGAATGGC |
| USH2A_E63_fw | ATAGTAAACCAGCTGAAGCC |
| USH2A_E71_fw | CTCATGGACATTCAAGACAAG |
| USH2A_E72_fw | CTCTGTCTAACACACAATAGT |
| USH2A_E7_rev | TTTCCGTTGGTTGTGGACTA |
| USH2A_E14_rev | TGGCATTGCCTGGAGAAATA |
| USH2A_E21_rev | TTCCTTTAACCAGAGGTGGC |
| USH2A_E30_rev | CAGGTCACCTCAATGCTGTAT |
| USH2A_E40_rev | ACTATGTGCACTGCCAAATCC |
| USH2A_E46_rev | GAGATGTCCAGATGACACGTA |
| USH2A_E55_rev | CTTGGGTAGTAGCTGCAACTA |
| USH2A_E63_rev | CCGTCACTGAAGATGTTGTAT |
| USH2A_E64_rev | CATCAAAGGTGCAATCTCAG |
| USH2A_E72_rev | GTTCATCAGGTCCTCTTCAT |
| USH2A_E72.1_rev | GACTATCCCTTACTTTACAAG |
| USH2A_E72.2_rev | GGTCACATTCAAATATTTATTAA |

**Suppl. Table 5: RT-qPCR primer**

| Primer | Sequence [5’ -> 3’] |
|---|---|
| qPCR_RPE65_fw | CAGCTCATGTAACAGGCAGGA |
| qPCR_RPE65_rev | GCTTGCCCATCAAACAGGTG |
| qPCR_ABCA4_fw | GGCCTCCACCACAAGCGGAATG |
| qPCR_ABCA4_rev | CAGGAGCAGATCCCAGATTGAGCGT |
| qPCR_MYO7A_fw | TGGTGATTCTTCAGCAGGGG |
| qPCR_MYO7A_rev | GGAGAGATCCAGTGTTCATTGTC |
| qPCR_USH2A_fw | CAGGCCAGTGCAAGTGCAAAGC |
| qPCR_USH2A_rev | ACTGGCAGGGCTCACATCCAAC |
